# Autofluorescence is a biomarker of neural stem cell activation state

**DOI:** 10.1101/2022.12.14.520430

**Authors:** Christopher S. Morrow, Kelsey Tweed, Zachary P. Arndt, Alex J. Walsh, Bo Peng, Ryan D. Risgaard, Payton C. Klosa, Michelle M. Chi, Eli P. Wallace, Mathew V. Jones, Avtar Roopra, Melissa C. Skala, Darcie L. Moore

## Abstract

Neural stem cells (NSCs) in the adult brain are primarily quiescent but can activate and enter the cell cycle to produce newborn neurons. NSC quiescence can be regulated by disease, injury, and age, however our understanding of NSC quiescence is limited by technical limitations imposed by the bias of markers used to isolate each population of NSCs and the lack of live-cell labeling strategies. Fluorescence lifetime imaging (FLIM) of autofluorescent metabolic cofactors has previously been used in other cell types to study shifts in cell states driven by metabolic remodeling that change the optical properties of these endogenous fluorophores. Here we asked whether autofluorescence could be used to discriminate NSC activation state. We found that quiescent NSCs (qNSCs) and activated NSCs (aNSCs) each have unique autofluorescence intensity and fluorescence lifetime profiles. Additionally, qNSCs specifically display an enrichment of a specific autofluorescent signal localizing to lysosomes that is highly predictive of cell state. These signals can be used as a graded marker of NSC quiescence to predict cell behavior and track the dynamics of quiescence exit at single cell resolution *in vitro* and *in vivo*. Through coupling autofluorescence imaging with single-cell RNA sequencing *in vitro* and *in vivo*, we provide a high-resolution resource revealing transcriptional features linked to rapid NSC activation and deep quiescence. Taken together, we describe a single-cell resolution, non-destructive, live-cell, label-free strategy for measuring NSC activation state *in vitro* and *in vivo* and use this tool to expand our understanding of adult neurogenesis.

## Introduction

Neural stem cells (NSCs) are responsible for the lifelong production of newborn neurons, a process referred to as neurogenesis, in at least two distinct zones: the subventricular zone of the lateral ventricles and the subgranular zone of the dentate gyrus in the hippocampus [1]. Adult NSCs are primarily quiescent (qNSCs), but upon receiving a signal will activate, exiting quiescence and entering into the cell cycle. These activated NSCs (aNSCs) can both self-renew and produce progenitors which expand the population, before ultimately differentiating into newborn neurons which mature and integrate into the existing circuitry [1]. Recent evidence suggests that NSC quiescence exit is the rate-limiting step in adult neurogenesis and is decreased with age and disease [2-5]. Thus, understanding the biology of qNSCs, aNSCs and the transition from quiescence to activation has become critical to understanding adult neurogenesis. Studies over the past decade have provided insight into the biology underlying NSC quiescence, revealing widespread remodeling of many nodes of cell biology, such as metabolism and the proteome [2, 6-12]. However, these studies have been limited by the constraints of modern technologies which exist to identify, isolate and/or generate qNSCs and aNSCs. For example, intermediate filaments, such as nestin and glial fibrillary acidic protein (GFAP) are commonly used as markers to isolate NSCs, yet these proteins are highly differentially expressed in qNSCs and aNSCs [6, 10, 11, 13, 14]. To gain the most complete understanding of NSC quiescence, new tools with unique capabilities need to be applied.

Many studies have demonstrated the potential of fluorescence lifetime imaging (FLIM) of autofluorescence to study shifts in cell state and behavior in other cell types [15-19]. FLIM measures the rate of signal decay of a fluorophore after excitation, called the fluorescence lifetime, which can be used to extrapolate biophysical properties of a fluorophore (Fig. 1A). For example, nicotinamide adenine dinucleotide phosphate (NAD(P)H; NADPH and NADH autofluorescence are indistinguishable and thus NAD(P)H is used to represent the combination) and flavin adenine dinucleotide (FAD) autofluorescence have been used to resolve spatial and temporal dynamics in macrophage metabolism, and to classify T-cell activation state [15, 16, 20]. Autofluorescence imaging capitalizes on the principle that as autofluorescent molecules are used by cells in different ways (e.g. binding to an enzyme, oxidation state) across cell states, identities or behaviors, their optical properties will change. For example, NAD(P)H is autofluorescent in the reduced state, whereas NAD(P) in the oxidized state is not autofluorescent [21]. Conversely, FAD is autofluorescent in the oxidized state, but not autofluorescent in the reduced state (FADH_2_) [15, 22]. In addition to detecting relative abundance of NAD(P)H or FAD through measuring their fluorescence intensity, FLIM can provide details on the biophysical properties of NAD(P)H and FAD. For example, NAD(P)H has a shorter fluorescence lifetime when it is not bound to a protein (Table 1) [17]. Conversely, FAD has a longer lifetime when it is not bound to a protein [21, 22]. NAD(P)H and FAD are known to be involved in hundreds of enzymatic reactions in the cell. Thus, it can be challenging to identify the precise underlying binding partners responsible for changing NAD(P)H and FAD fluorescence lifetimes across conditions. However, regardless of what is specifically driving changes in their fluorescence lifetimes, measuring differences in NAD(P)H and FAD autofluorescent intensity and lifetimes can be sufficient on its own to inform on a cell’s overall behavior. Thus, we asked whether a combination of lifetime and intensity imaging of specific autofluorescent signals (referred to here as optical cell state imaging; OCSI) could be used to study NSC cell state. We found that OCSI could be used as a non-destructive, single-cell resolution, graded, live-cell, label-free marker to classify NSC activation state. Using autofluorescence imaging combined with single cell RNA sequencing of NSCs *in vitro* and *in vivo*, we provide resources that reveal the molecular features associated with rapid quiescence exit and deep quiescence. This strategy provides a unique, easily accessible platform to pair a functional readout of NSC metabolism with gene expression, expanding our understanding of NSC quiescence.

**Table 1.**
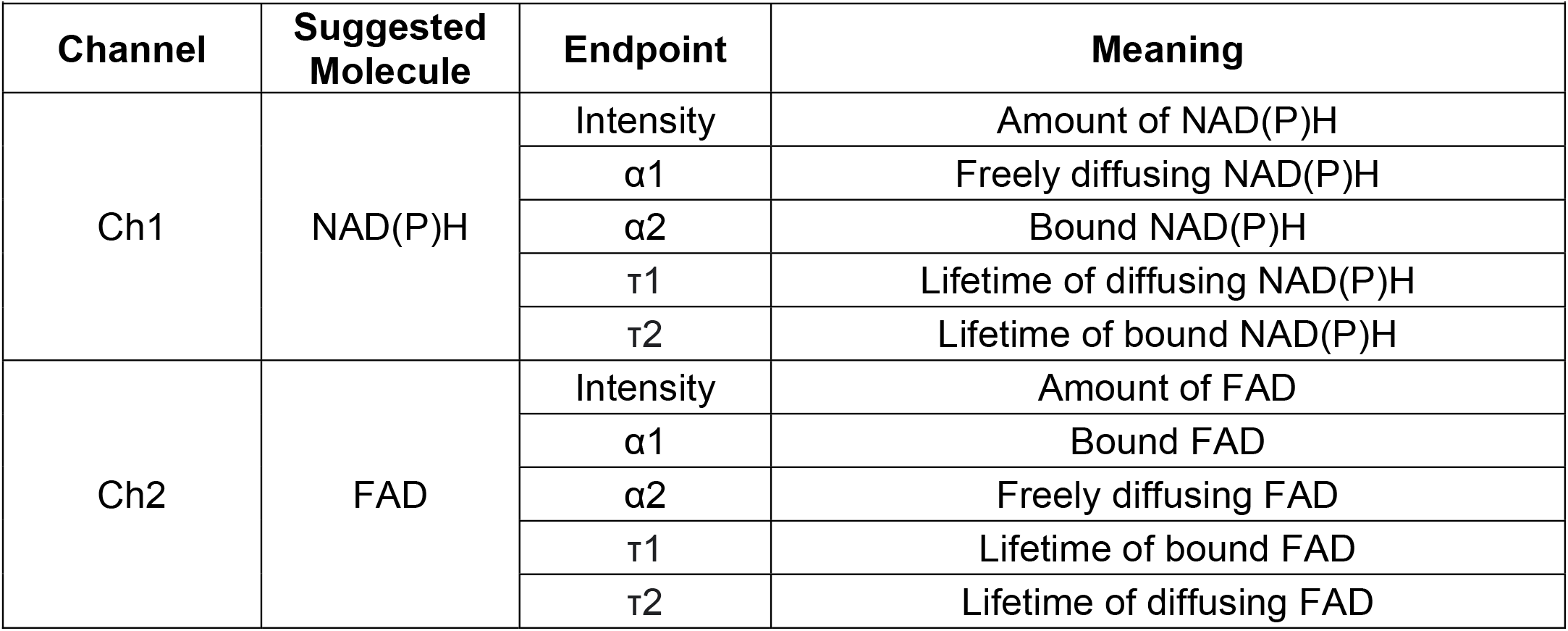
Autofluorescent endpoints.

**Figure 1.**
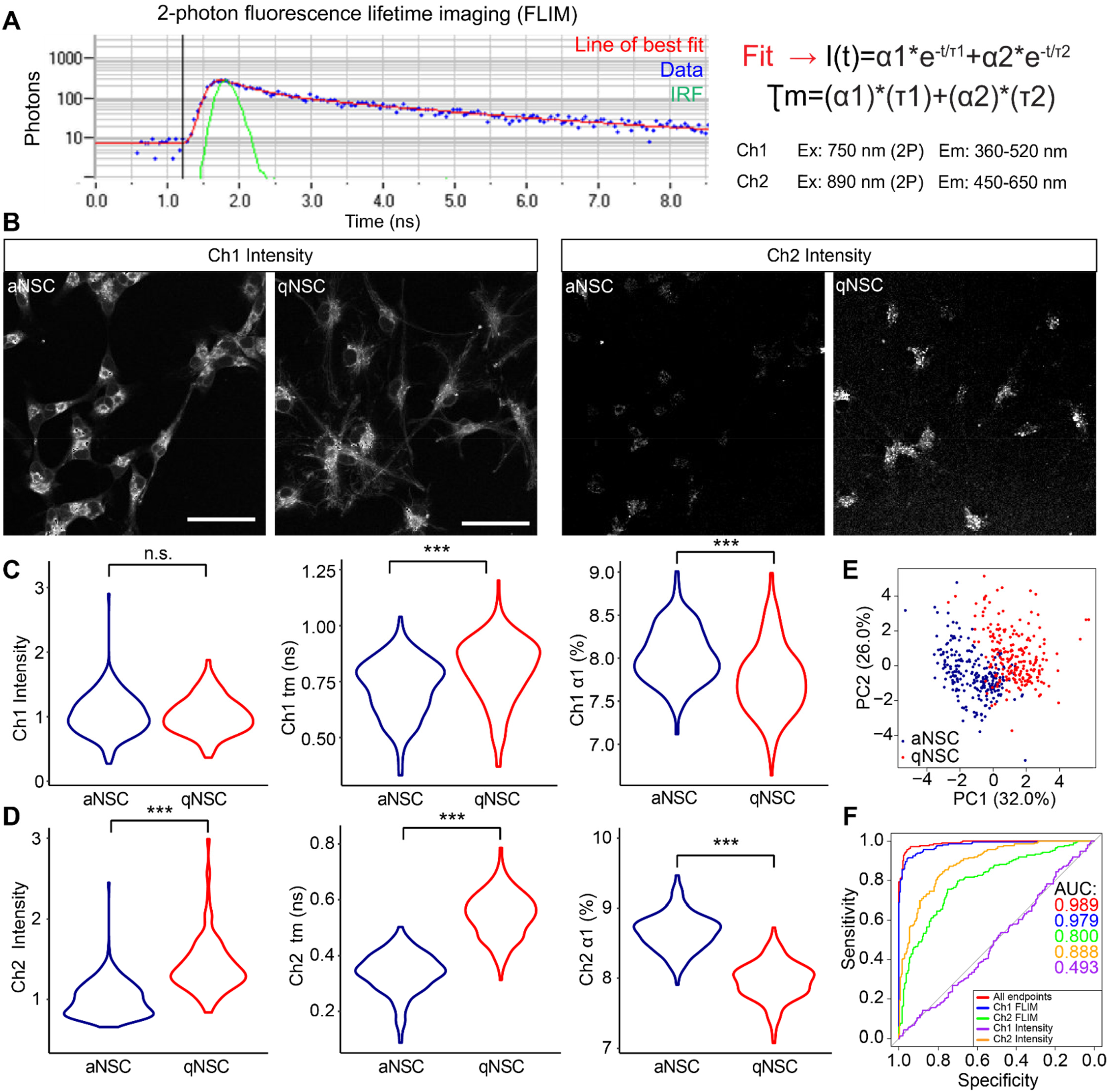
A live-cell, label-free imaging strategy for the classification of NSC activation state. A) Schematic depicting fluorescence lifetime imaging (FLIM) analysis. Data (blue) is modeled by a biexponential decay equation (red). The instrument response function (IRF) is shown in green. B-D) aNSCs and qNSCs were imaged for Ch1 and Ch2 intensities and fluorescence lifetimes. (B) 2-photon intensity images of Ch1 and Ch2 in qNSCs and aNSCs. (C-D) Violin plots depicting intensity and representative FLIM endpoints for Ch1 and Ch2 in qNSCs (blue) and aNSCs (red) (N=3, Generalized Linear Model). E) Principle component analysis of qNSC (red) and aNSC (blue) OCSI data (Ch1 and Ch2 intensity, α_1_, τ_1_, and τ_2_). F) Receiver operating characteristic curve depicting a random forest model generated to classify NSC activation state using NSC autofluorescence data. Different lines represent random forest models constructed using subsets of NSC autofluorescence data. Scale bars, 50 μm. ***p < 0.001.

## Results

### Autofluorescence can be used to accurately classify NSC activation state

To determine whether autofluorescence imaging could be used to classify NSC activation state, we first isolated primary hippocampal NSCs from 6 week-old mice and generated qNSCs and aNSCs *in vitro* using a previously established paradigm using bone morphogenic protein 4 (BMP4) to drive aNSCs into quiescence (Fig. S1A-B) [6, 7, 13, 23-25]. Using a 2-photon microscope, we performed FLIM on qNSCs and aNSCs *in vitro* using optical parameters designed to image the metabolic cofactors NAD(P)H (Channel 1 (Ch1) Excitation: 2-photon 750 nm, Emission: 360-520 nm) and FAD (Channel 2 (Ch2) Excitation: 2-photon 890 nm, Emission: 450-650 nm) (Fig. 1A) [26]. qNSCs and aNSCs on average exhibited significantly different Ch1 and Ch2 OCSI endpoints (Table 1; Fig. 1B-D). Principal component analysis (PCA) of qNSCs and aNSCs using all 8 OCSI measures (Intensity, α1, τ1 and τ2 for both Ch1 and Ch2) revealed clear separation between qNSCs and aNSCs, suggesting the predictive power of autofluorescence to assess NSC cell state (Table 1, Fig. 1E). To further evaluate whether NSC autofluorescence could be used to predict NSC activation state, we constructed classification models using random forest machine learning and varying combinations of autofluorescent endpoint inputs (Fig. 1F). Receiver operator characteristic (ROC) curves illustrated that whereas training the model with only Ch1 intensity (purple line) resulted in poor classification no greater than randomly guessing whether a NSC was quiescent or activated, training the model with all 8 endpoints (Intensity, α1, τ1 and τ2 for both Ch1 and Ch2; red line) resulted in an almost perfect model. Subsets of autofluorescent endpoints, such as Ch1 FLIM, also revealed strong modeling. Strikingly, we observed that a single endpoint, Ch2 intensity, alone was sufficient to provide robust classification of NSC activation state (Fig. 1F). Taken together, these data demonstrate distinct differences in autofluorescence properties between qNSCs and aNSCs and highlight the potential of autofluorescence to be used as a biomarker of NSC activation state.

### The autofluorescent signal detected by Ch2 is in lysosomes, and is highly enriched in qNSCs

As expected from previous studies, the autofluorescence signal in Ch1 attributed to NAD(P)H localized to mitochondria (Fig. 1B, Table 2) [27]. However, despite using excitation and emission spectra consistent with published reports to excite and detect FAD (Ch2), we observed punctate signals in qNSCs not reminiscent of FAD, which others had shown to be diffusely spread throughout the cytoplasm (Fig. 1B) [15]. To gain further insight into the optical properties of these signals, we first used both confocal microscopy and fluorescent activated cell sorting (FACS) to identify where the puncta would absorb and emit light. Interestingly, we observed that autofluorescent puncta in qNSCs were broadly excited by 405-647 nm light and emitted between 500-700 nm (Fig. S1C-D). These data further suggest that the puncta are not FAD, as FAD previously has not been shown to be highly excited by a 561 nm laser [28]. Next, we measured NAD(P)H and FAD levels in cell lysates, and found that both qNSCs and aNSCs contained similar levels of both molecules, suggesting that varying levels of these factors likely were not contributing to differences in autofluorescence intensity observed across aNSCs and qNSCs (Fig. S2A-B). Finally, if FAD were substantially contributing to the intensity of the Ch2 signal, this signal should be sensitive to treatment with Carbonyl cyanide-p-trifluoromethoxyphenylhydrazone (FCCP), which should make FAD more autofluorescent, or sodium cyanide (NaCN), which should make FAD less autofluorescent [29-31]. Whereas Ch1 autofluorescence in qNSCs and aNSCs was sensitive to NaCN and FCCP as expected if detecting NAD(P)H and consistent with previous reports, treatment with FCCP and NaCN showed no effect on the Ch2 signal in qNSCs, further confirming that FAD is not substantially contributing to the signal detected using Ch2 (Fig. S2C-F). Thus, we hereafter call the autofluorescent signal we detect with Ch1 NAD(P)H, and conservatively describe the autofluorescent signal detected with Ch2 **P**unctate **A**uto**f**luorescence (PAF; Fig. S2G).

**Table 2.** OCSI Imaging Strategies.

| **Table 2 - OCSI Imaging Strategies** |  |  |  |  |
| --- | --- | --- | --- | --- |
| **Channel Name** | **Detection System** | **Excitation (nm)** | **Emission (nm)** | **Fluorescent Species Detected** |
| Ch1 | 2-photon | 750 (375 – 1P) | 360-520 | NAD(P)H |
| Ch2 | 2-photon | 890 (445 – 1P) | 450-650 | PAF |
| N/A | Confocal | 405 | 525-640 nm | PAF |
| N/A | Flow Cytometer | 405 | 585-625 | PAF |

Further examination of the morphology of the PAF enriched in qNSCs revealed the signal was enriched in round structures. As qNSCs and aNSCs have distinct differences in protein homeostasis, and qNSCs are marked by an accumulation of lysosomes, we hypothesized that the PAF may be in lysosomes [6, 7, 9]. To test this, we labeled qNSCs and aNSCs with a live, lysosome-specific dye (Lysotracker far red to avoid bleed-through from the dye) and performed autofluorescence intensity imaging using a confocal microscope to detect the PAF (Excitation: 405 nm laser, Emission: 525-560 nm). The PAF exhibited 97.98±0.01% overlap with the Lysotracker dye, suggesting that these punctate signals present in qNSCs are associated with lysosomes (Fig. 2A-E, Table 2). However, most lysosomes were clearly not autofluorescent (only 29.82±0.09% overlap), suggesting that the PAF signal is specifically labeling a subpopulation of lysosomes in qNSCs (Fig. 2D).

**Figure 2.**
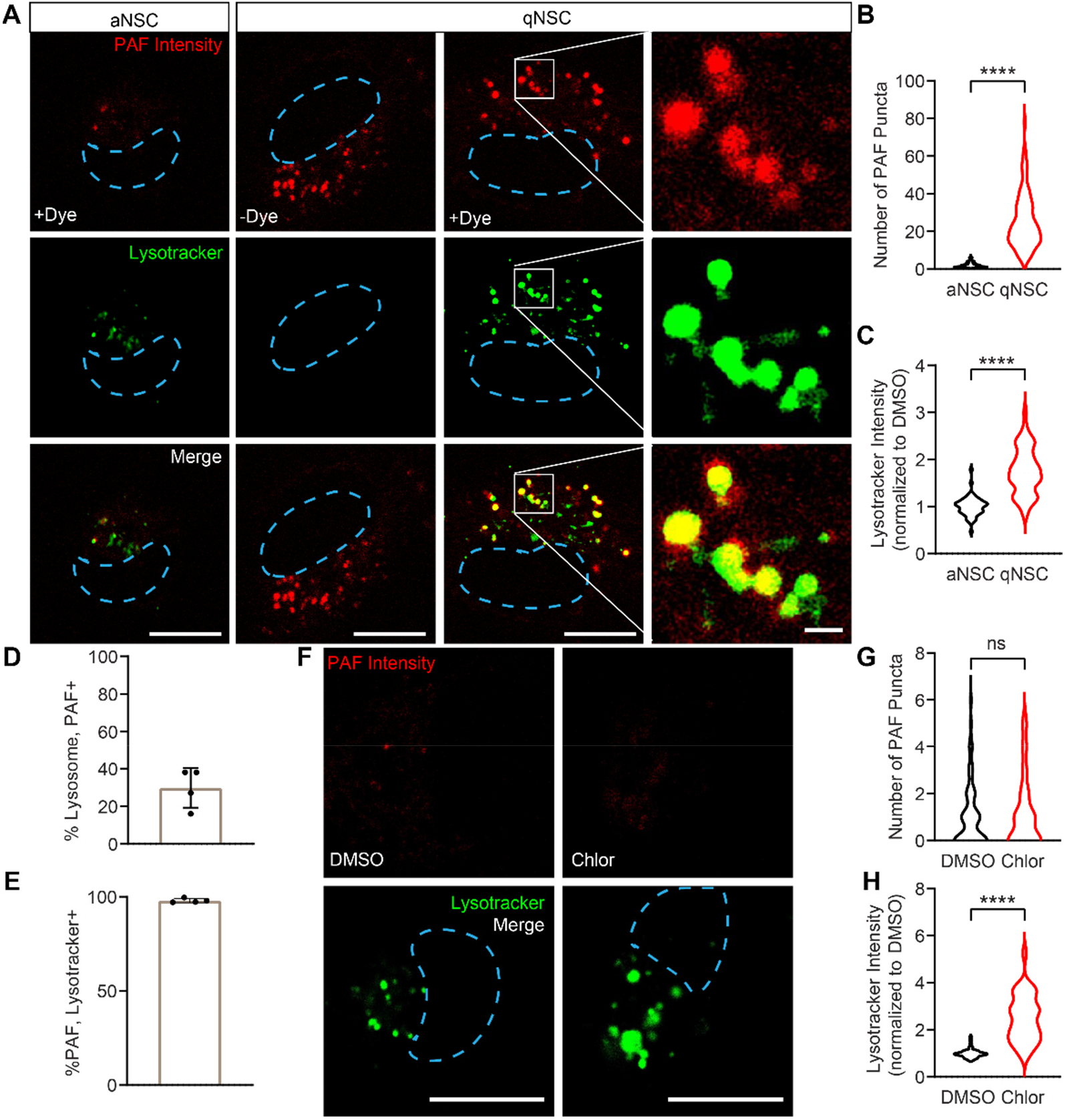
Autofluorescence localizing to lysosomes in qNSCs marks NSC activation state. A-E) qNSCs and aNSCs were labeled with Lysotracker and then imaged on a confocal microscope and analyzed for lysosomes (Confocal, green, Ex: 405 nm Em: 525-560 nm) and PAF, and analyzed for the abundance of Lysotracker fluorescence intensity, number of PAF (Confocal; red, Ex: 405 nm Em: 580-620 nm), and %colocalization between PAF and Lysotracker (N=3, Student’s t test, mean ± SD). F-H) aNSCs were treated with either DMSO or 10 μM chloroquine for 2.5 hours, stained with Lysotracker to label lysosomes, and then imaged with a confocal microscope and analyzed for PAF (red) and lysosomes (green) (N=3, Student’s t test, mean ± SD). Nuclei are outlined by a blue dotted line. White line denotes inset. Scale bars, 10 μm (outsets), 1 μm (insets). ****p < 0.0001.

We next asked if increasing lysosomes was sufficient to increase PAF levels. Previously it has been shown that chloroquine treatment rapidly induces the accumulation of lysosomes [32]. Thus, we treated aNSCs with 10 μM chloroquine or DMSO for 2.5 hours and found that although NSCs did exhibit higher levels of lysosomes, the PAF signal did not similarly increase (Fig. 2F-H), suggesting that increasing lysosomes was not sufficient to increase PAF. Thus, these data collectively suggest that qNSCs exhibit high levels of an autofluorescent signal unique to a subpopulation of lysosomes.

### Autofluorescence can predict NSC activation state and proliferative behavior in a mixed population *in vitro*

Could NSC autofluorescence be used to predict NSC activation state and cell behavior in a mixed population of qNSCs and aNSCs? As we had previously observed that qNSCs are marked by higher levels of PAF, we first asked whether using FACS to sort NSCs based on PAF intensity would be sufficient to enrich separately for qNSCs and aNSCs. To this end, we generated qNSCs and aNSCs *in vitro*, labeled both groups with a 1 hour EdU pulse, and then sorted cultured qNSCs only, cultured aNSCs only, and a 1:1 mixture (Mix) of these two populations based on their PAF intensity (FACS: 405 Excitation, 560-640 Emission) (Fig. 3A-B). Confirming what we observed in plated cells *in vitro*, qNSCs had higher PAF intensity than aNSCs using this FACS-based imaging strategy (Fig. 3B). We then sorted the Mix cells using gates drawn to separately collect NSCs with brighter (High) or dimmer (Low) PAF intensity and plated them, allowing them to adhere to the dish over the course of 2-3 hours. Sorted cells were fixed, immunostained, and analyzed for EdU to mark cell cycle status in each population of cells (Fig. 3C-D). Whereas the Mix cells exhibited a proliferation rate in between qNSCs and aNSCs as expected, High PAF cells purified from the Mix proliferated more similarly to qNSCs and Low PAF cells purified from the Mix proliferated more similarly to aNSCs. Thus, FACS based solely on PAF intensity can be used to predict NSC activation state and proliferative behavior.

**Figure 3.**
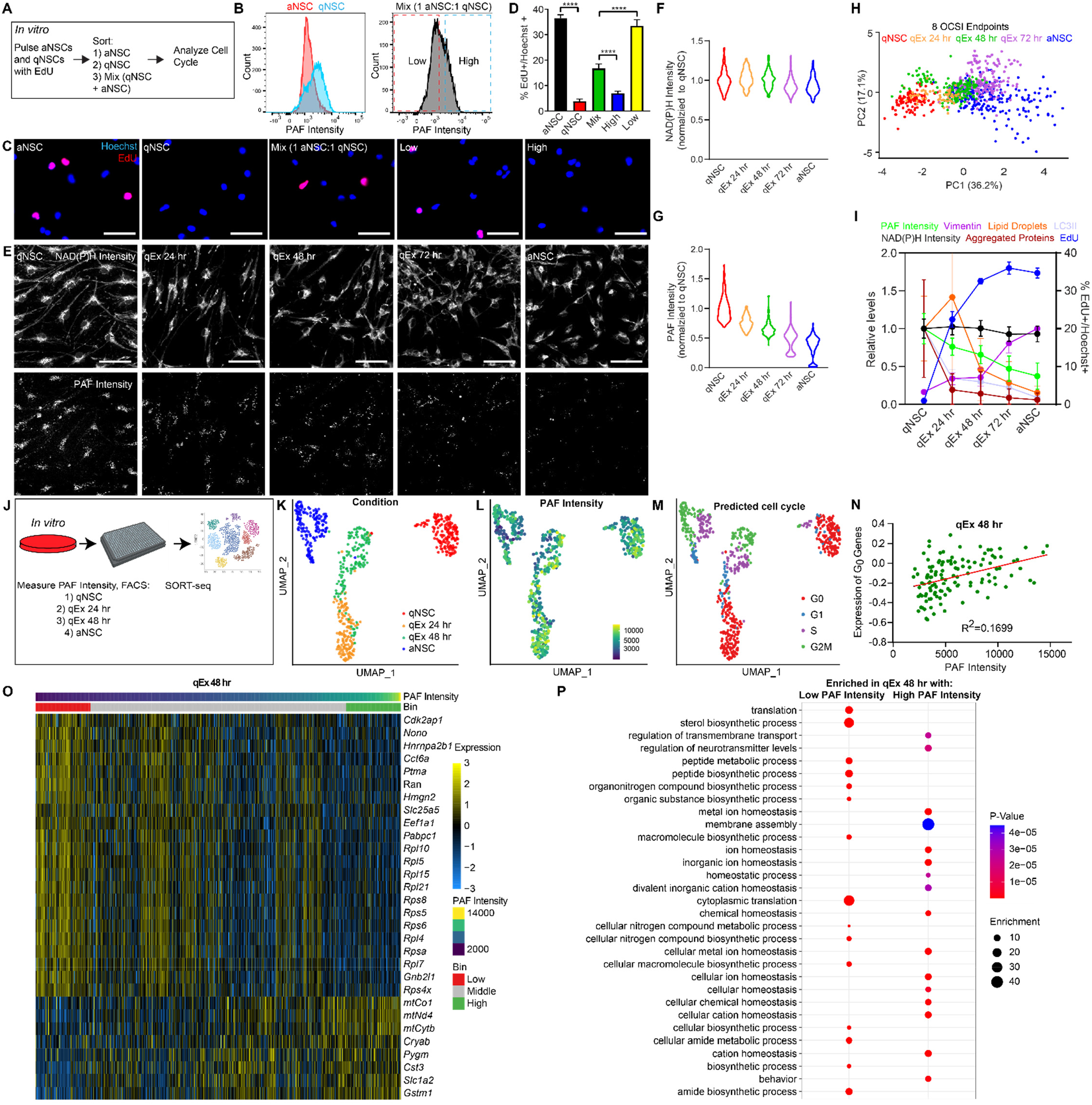
NSC autofluorescence can enrich for NSC activation state and be used to identify transcriptional signatures of rapid NSC activation. A-D) qNSCs and aNSCs were pulsed with EdU for 1 hour and then prepared for cell sorting either separately or in a 1 aNSC:1 qNSC mixture and then sorted based on relatively low or high PAF intensity (Ex: 405 nm Em: 580-620 nm). 3 hours after plating, cells were fixed, treated to visualize EdU (red) and nuclei (Hoechst; blue) and analyzed for %EdU+ of total Hoechst+ cells (N=3, Student’s t test, mean ± SD). E) NAD(P)H (top) and PAF (bottom) 2-photon intensity images of a quiescence exit time course. F-G) Analysis of NAD(P)H and PAF intensity of the quiescence exit time course (n=783 cells). H) PCA of *in vitro* NSC quiescence exit autofluorescence imaging data (NAD(P)H and PAF intensity, α_1_, τ_1_, and τ_2_). I) Overlay of changes in autofluorescence intensity during the quiescence exit time course combined with the proliferation rate marker (EdU; blue), LC3II (light blue), aggregated proteins (Proteostat; red), vimentin (purple) and lipid droplets (nile red; orange). J) *In vitro* qNSCs, aNSCs, and qNSCs exiting quiescence for either 24 (qEx 24 hr) or 48 (qEx 48 hr) hours were analyzed by FACS to measure PAF intensity, single-cell sorted, and sent for single-cell RNA sequencing (SORT-seq). K-N) UMAP analysis of *in vitro* SORT-seq cell transcriptomes labeled by sample condition (K), PAF intensity (L), and predicted cell cycle stage (M). N) Plot comparing PAF intensity to expression of G_0_ genes in qNSCs exiting quiescence for 48 hours (qEx 48 hr) from *in vitro* SORT-seq [36]. O) Heat map displaying normalized expression of the top 30 differentially expressed genes when comparing the top 15% to the bottom 15% PAF intensity qEx 48 hr cell transcriptomes from *in vitro* SORT-seq. P) Dot plot depicting the top 30 gene ontology terms enriched in either the top 15% or bottom 15% PAF qEx 48 hr cell transcriptomes from *in vitro* SORT-seq. Scale bars, 50 μm. ****p < 0.0001.

### Autofluorescence imaging reveals the heterogeneity in the rate of quiescence exit *in vitro*

As NSC autofluorescence can classify NSC activation state (Fig. 1, 3A-D), we next hypothesized that autofluorescence could provide a window into the dynamics of NSC quiescence exit and act as an additional sensor to monitor cell behavior during NSC quiescence exit. To test this hypothesis, we used a 2-photon microscope to perform OCSI in NSCs at fixed timepoints as qNSCs exited quiescence over the course of three days until fully activated (Fig. 3E-H). As expected, we observed minimal changes in NAD(P)H autofluorescence intensity as NSCs exited quiescence, whereas at the same time, PAF intensity steadily decreased towards levels observed in aNSCs (Fig. 3F-G). These data suggest that PAF intensity could act as a graded marker of NSC activation state at the population level.

To determine if OCSI at the single-cell level could be used to detect the heterogeneity of *in vitro* quiescent exit through dimension reduction analysis, we performed OCSI at various stages of quiescence exit, including quiescence, 24, 48, and 72 hours following addition of activation media, and fully activated (aNSCs). Interestingly, using only OCSI data, PCA revealed a step-wise trajectory of quiescence exit, and the single-cell heterogeneity in the rate of quiescence exit (Fig. 3H). For example, cultures of qNSCs that had been treated with activation media for 24 hours contained cells that clustered within the group of qNSCs, suggesting that these cells had not begun to exit quiescence yet, while some cells had moved into an intermediate state where they did not cluster with qNSCs or aNSCs. Conversely, cultures of qNSCs that had been treated with activation media for 48 hours had cells that clustered with aNSCs that had more fully reactivated in addition to cells that clustered in an intermediate state. Importantly, the time from quiescence exit was reflected in a clear trajectory towards aNSCs, revealing the strength of autofluorescence imaging to report at single-cell resolution measures of functional, dynamic cellular changes occurring during shifts in NSC cell state. Thus, OCSI provides a novel strategy to identify distinct substates of NSCs and track the dynamics of NSC quiescence exit *in vitro*.

To determine if the shifts in autofluorescence signals during quiescence exit are dynamically similar to changes occurring in other reported markers, we compared NAD(P)H and PAF intensity to other markers that are known to shift during NSC quiescence exit: 1) aggregated proteins labeled by the dye Proteostat, 2) EdU to indicate proliferation rate, 3) vimentin protein levels, 4) lipid droplets labeled by Nile red, and 5) the levels of LC3II indicative of autophagic flux (Fig. 3I, S3A-D) [6, 13, 33, 34]. Previously it was shown that as NSCs exit quiescence, aggregated proteins decrease while proliferation markers and vimentin protein levels increase [6, 13]. Thus, at the same time points in which we performed autofluorescence imaging, we analyzed levels of each of these markers as well as a marker of autophagic flux. Expectedly, we observed that as NSCs exited quiescence, they became more proliferative, cleared the Proteostat+ aggregated proteins, had decreased levels of LC3II, degraded lipid droplets, and increased vimentin protein levels (Fig. 3I). Interestingly, we observed that EdU and Proteostat had the most rapid changes during quiescence exit, whereas PAF intensity and vimentin protein levels changed more gradually as NSCs progressed towards activation. This suggests that PAF intensity may act as a more dynamic marker of quiescence exit, providing greater resolution of distinct substages throughout quiescence exit. Altogether, these data provide a proof-of-principle for how autofluorescence signals can be used to study NSC quiescence, both through using autofluorescence to predict NSC activation state and cell behavior, as well as providing greater resolution of stages of quiescence exit and their underlying dynamics.

### PAF intensity coupled with gene expression during NSC quiescence exit *in vitro* reveals features associated with rapid NSC activation

We next capitalized on the power of PAF intensity to predict the heterogeneity in NSC quiescence exit *in vitro* and used it to identify transcriptional features associated with cells that exited quiescence more or less quickly. We generated qNSCs, aNSCs, and NSCs exiting quiescence for 24 or 48 hours *in vitro*, sorted single cells using FACS into 384 well plates, tracking PAF intensity for each cell, and then performed single cell RNA sequencing on the resulting cells (Fig. 3J, S3E-G, Table S1; SORT-seq, Single Cell Discoveries) [35]. For each condition, we observed the spread of PAF intensity, made 4 distinct gates within that spread, and collected equal numbers of cells from each gate to obtain a representation of the whole condition’s population (Fig. S3E). Uniform Manifold Approximation and Projection (UMAP) analysis demonstrated strong separation of each sample based on condition, with cells progressively moving between quiescence and activation (Fig. 3K, S3H-I). In agreement with our previous data, qNSCs had the brightest PAF intensity, and PAF intensity progressively decreased as qNSCs exited quiescence in our 24 and 48 hour timepoints (Fig. 3L).

As we sampled a range of PAF intensity signals per condition, we hypothesized that variance in PAF intensity could be connected to heterogeneity in the rate of quiescence exit and/or activation state in culture. To test this hypothesis, we measured the expression of G_0_ genes [36], and examined these in relation to PAF (Fig. 3M, S3J). We observed in our 48 hour exiting quiescent samples that higher PAF intensity was associated with higher expression of G_0_ genes (Fig. S3N), suggesting that PAF intensity could be used to identify the features of rapid NSC activation. To identify these features, we performed a differential expression analysis comparing the top 15% PAF intensity cells to the bottom 15% PAF intensity of cells in our 48 hour quiescence exit condition (Fig. 3O-P, S3K-L). Using a log transformed fold change of 0.75 or greater as a cutoff, we identified 35 genes that were more highly expressed in high PAF cells and 98 genes which were more highly expressed in low PAF cells (Table S2). Gene ontology analysis of these terms revealed nodes of genes enriched in high or low PAF cells, displaying agreement with previous studies demonstrating distinct transcriptional differences between aNSCs and qNSCs [6, 10-12]. For example, mRNA translation has been reported to be increased in aNSCs and aNSCs and qNSCs are known to substantially remodel their metabolism. Indeed, in low PAF intensity cells we observed upregulation of many genes linked to translation and metabolism of many different types of molecules, such as peptides and amides (Fig. 3O-P, Table S3). These findings demonstrate that the live measure of PAF intensity is predictive not only of NSC activation state at the level of cell behavior (Fig. 3A-H), but also at the transcriptional level (Fig. 3K-P). Thus, PAF can reliably report on NSC cell state without the need for destruction of the cell nor any additional labels, and can be used to identify features linked to rapid NSC activation at the single cell level.

### PAF are detectable in qNSCs in the mouse brain

Our *in vitro* observation of PAF enrichment in qNSCs raised the prospect that autofluorescence could be used to study NSC quiescence in the brain. To address whether autofluorescent features observed *in vitro* in qNSCs and aNSCs are conserved in the mouse brain, we stereotactically injected adeno-associated virus serotype 4 (AAV4) particles encoding a CMV promoter driving expression of cytosolic mCherry, a fluorophore compatible with our 2-photon FLIM strategy, into the hippocampus [37]. Seven days following injection, to allow for detectable expression of the viral reporter in live slices, we generated acute organotypic slice cultures to image PAF intensity in the mouse hippocampus [38] (Fig. 4A). Previous studies have suggested AAV4 preferentially transduces qNSCs when injected into the hippocampus [37], thus we first validated AAV4-CMV-mCherry labeling of qNSCs in our own hands. We found that mCherry+ cells in the SGZ of the dentate gyrus of the hippocampus were 94.4±1.79% Sox2+/BrdU-, confirming previous findings [37] (Fig. S4A). Thus, using this system, we performed autofluorescence imaging on qNSCs in the mouse brain using 2-photon imaging to visualize the autofluorescence intensity of NAD(P)H and PAF in mCherry+ NSCs. While imaging PAF, we observed the presence of punctate autofluorescent signals reminiscent of our *in vitro* findings in qNSCs in mCherry+ qNSCs in the mouse hippocampus (Fig. 4A). This suggested that NSC autofluorescence may be used *in vivo* to mark NSC activation state.

**Figure 4.**
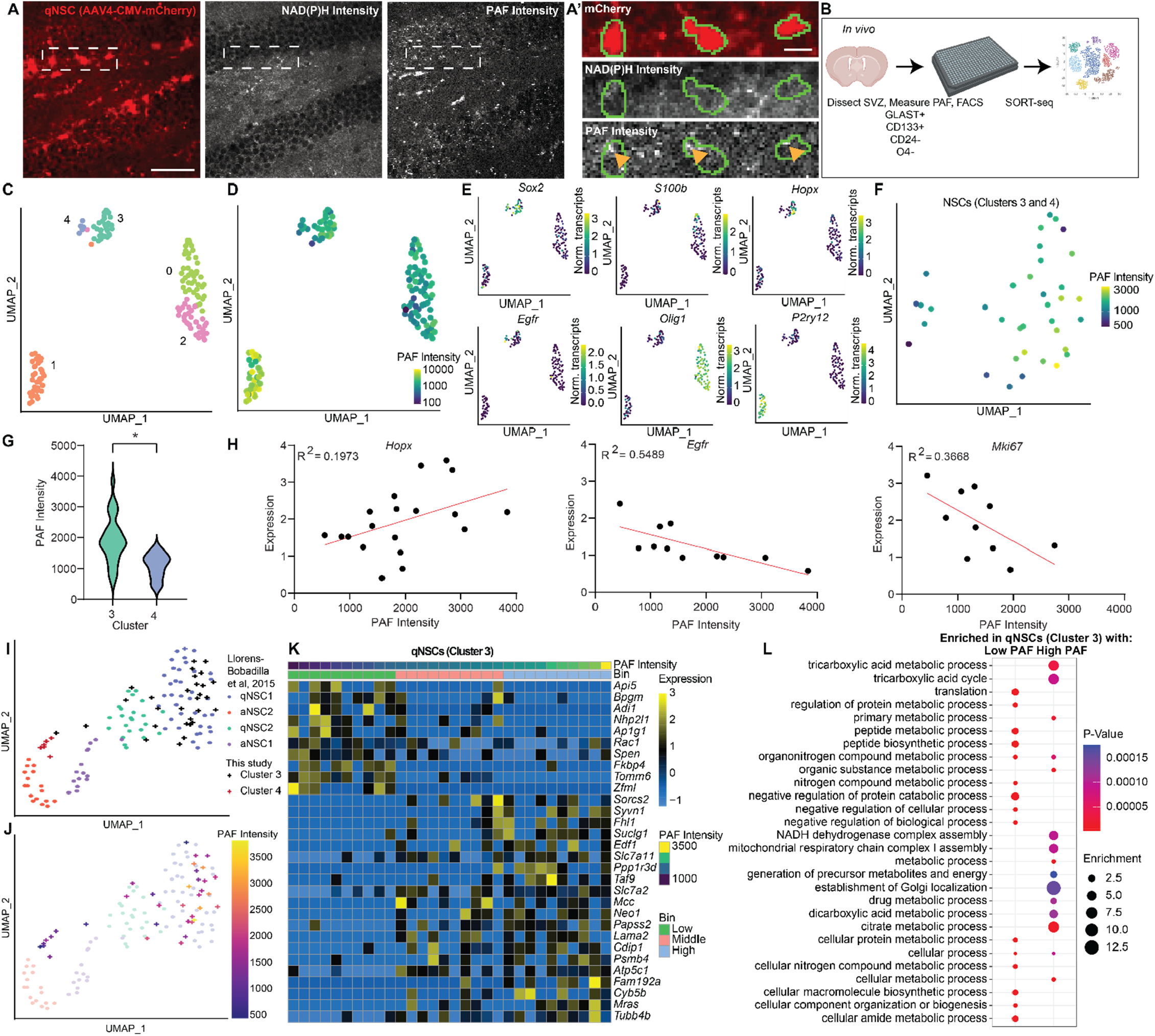
PAF are detectable in qNSCs in the mouse brain and are a graded marker of deep quiescence. A) Mice were stereotactically injected with AAV4-CMV-mCherry viral particles into the dentate gyrus of the hippocampus, and allowed to recover for 7 days. Brains were extracted and live, organotypic slice-cultures were prepared. Slices were immediately imaged with a 2-photon microscope for mCherry (red), NAD(P)H (white), and PAF (white). Green lines denote the edge of qNSCs. Yellow arrows denote autofluorescent puncta. B) Schematic depicting strategy for enriching for NSCs isolated from the SVZ, tracking PAF intensity by FACS, and sending for SORT-Seq. C) UMAP analysis of *in vivo* SORT-seq transcriptomes labeled by clusters identified using the Louvain method. D) UMAP analysis of single cell transcriptomes from cells isolated from the SVZ as described in Fig. 4B with PAF intensity annotated. E) UMAP analysis of single cell transcriptomes from cells isolated from the SVZ as described in Fig. 4B with expression of markers of different cell types and cell states annotated. F) UMAP analysis of single cell transcriptomes of NSC harboring clusters (clusters 3 and 4) isolated from the SVZ as described in Fig. 4B with PAF intensity annotated. G) PAF Intensity of cells in cluster 3 and 4 from *in vivo* SORT-seq (n=36 cells, Student’s t test, mean ± SD). H) Plots of *Hopx, Egfr* and *Mki67* expression as a function of PAF intensity, excluding cells with an expression of 0 for each gene respectively from *in vivo* SORT-seq. Trend lines are shown in red. I-J) UMAP analysis of NSC transcriptomes reported in Llorens-Bobadilla et al. 2015 (circles) overlayed with NSCs from clusters 3 and 4 produced by this study and J) PAF intensity (plus symbols; *in vivo* SORT-seq). K) Heat map showing normalized expression of the top 30 most differentially expressed genes comparing the top third to the bottom third PAF intensity of qNSCs (cluster 3) from *in vivo* SORT-seq. L) Dot plot depicting the top 30 gene ontology terms enriched in either the top third or bottom third PAF intensity qNSCs (cluster 3) from *in vivo* SORT-seq. Scale bars, 50 μm (A) 5 μm (A’). *p < 0.05.

### PAF coupled with gene expression of *in vivo* NSCs reveals features linked to deep quiescence

Visualization of PAF in qNSCs in live brain sections suggested we could purify NSCs from the adult brain and use FACS to couple PAF intensity with single cell RNA sequencing (SORT-seq) to study the underlying gene expression of each cell with respect to PAF across NSC quiescence *in vivo* [35]. To obtain a larger number of NSCs, we dissected the SVZ in adult mice, dissociated the tissue, and enriched for NSCs by isolating GLAST^+^/CD133^low-mid^/CD24^-^/O4^-^ cells by FACS while obtaining the PAF intensity for each cell [2, 10, 39, 40] (Fig. 4B, S4B, Table S4). Cells were plated into 384 wells and sent for SORT-Seq processing and analysis. UMAP analysis revealed several transcriptionally distinct clusters of cells with a wide range of PAF intensities (Fig. 4C-D). Using previously published markers of cell types in the SVZ, we identified qNSCs (cluster 3; Sox2+/S100β-/Egfr-), aNSCs (cluster 4; Sox2+/S100β-/Egfr+), oligodendrocytes (clusters 0 and 2; Olig1+) and microglia (cluster 1; P2ry12+) in the sequenced samples (Fig. 4E). Interestingly, microglia had the highest PAF signal of all cell types identified (Fig. 4D, S4C-D), supporting previous studies suggesting a subpopulation of microglia accumulate an autofluorescent signal derived from lysosomal storage bodies [41]. Upon removing non-NSC clusters, we next asked whether higher PAF intensity was associated with NSC quiescence *in vivo*. Excitingly, as we found with qNSCs *in vitro*, qNSCs *in vivo* exhibited higher PAF intensities relative to aNSCs (Fig. 4F-G). This finding highlights the capacity of autofluorescence to be used to discriminate qNSCs and aNSCs *in vivo*.

As we had found earlier that autofluorescence in NSCs *in vitro* could be used to track the dynamics of NSC quiescence exit, we hypothesized that NSC autofluorescence could be used as a graded marker to measure the dynamics of NSC quiescence exit *in vivo*, for prospective identification of cells in a transitioning state between quiescence and activation, or cells that are in “deeper” states of quiescence, as has previously been suggested [5, 10, 42]. To evaluate this hypothesis, we first plotted genes known to increase (*Mki67, Egfr*) or decrease (*Hopx*) with NSC quiescence exit against PAF intensity (Fig. 4H, S4E-F). Across all three markers, we observed the highest or lowest levels of PAF intensity to be associated with the highest or lowest expression of each marker respectively in a linear relationship. To further test this hypothesis, we analyzed our NSC single-cell transcriptomic data overlayed with single-cell transcriptomic data of SVZ NSCs from Llorens-Bobadilla et al 2015 [10] which had previously identified two qNSC substates (termed qNSC1 and qNSC2) in which qNSC2 cells were closer to activation (Fig. 4I-J). Excitingly, we observed that cells with the highest PAF intensity most closely resembled qNSC1 cells, with decreasing PAF levels as substates progressed to aNSC2, suggesting that PAF intensity could identify cells in a deeper state of quiescence, and act as a graded indicator of cell state.

To expand our understanding of the molecular features of deep quiescence, we performed a differential gene expression analysis of the top third PAF intensity qNSCs (cluster 3) vs. the bottom third PAF intensity qNSCs (cluster 3) (Fig. 4K-L, S4G). Using a log transformed fold change of 0.75 or greater as a cutoff, we identified 620 genes that were more highly expressed in high PAF qNSCs and 942 genes that were more highly expressed in low PAF qNSCs (Table S5). Within these lists, we identified genes which previously have been implicated as markers of NSC quiescence, such as Hopx which was found to be more highly expressed in high PAF qNSCs. To identify the biological processes which are regulated by the genes we found to be the most differentially expressed between high and low PAF qNSCs we performed a gene ontology analysis on these same lists of genes (Fig. 4M, Table S6). Our gene ontology analyses revealed many pathways which are enriched in either high or low PAF intensity qNSCs respectively. For example, we identified an enrichment of translation-related genes which were more highly expressed in low PAF intensity qNSCs, further supporting our observations from our *in vitro* single-cell RNA sequencing data sets and observations made by many others about translation changes that occur across NSC activation states [6, 10, 11]. Analogous to our *in vitro* SORT-seq data set, in this analysis we also identified many differentially regulated nodes of cellular metabolism, such as mitochondrial respiratory chain complex I assembly, which could reflect metabolic regulation of NSC quiescence which has been previously reported [7, 9, 25, 43]. Cumulatively, these analyses demonstrate that PAF intensity levels of acutely purified NSCs are indicative of cell state, even to the level of qNSC substates of resting (qNSC2) or deep (qNSC1) quiescence.

Together, this work demonstrates the power of autofluorescence to be used to study dynamic and heterogeneous processes at the single cell level. Specifically, we provide a graded, live-cell, non-destructive, label-free, single-cell resolution toolkit for studying NSC quiescence both *in vitro* and *in vivo* and use this tool to expand our understanding of NSC quiescence by identifying features linked to rapid activation and deep quiescence.

## Discussion

Autofluorescence provides an unprecedented tool as a live reporter of molecular processes, to understand the biology of NSCs and the dynamics underlying their changes in cell state as they progress towards generating newborn neurons. Using autofluorescence to identify NSC cell states and behaviors may offer significant technical advantages (e.g. live-cell, label-free, non-destructive, single-cell resolution, graded) without the biases and disadvantages of other canonical markers. For example, intermediate filaments such as nestin, vimentin, and glial fibrillary acidic protein (GFAP) are often used to identify NSCs in the brain in combination with other markers, however, these proteins are highly differentially expressed between qNSCs and aNSCs [6, 10, 11, 13, 14]. Therefore, studies relying on these markers may be missing important subpopulations or substates of NSCs in their analyses. Studies of NSCs using various NSC-specific promoters have further suggested that these lines each have distinct behaviors *in vivo* [4, 44, 45]. Thus, identifying additional tools that aren’t reliant solely on promoter-driven expression of proteins could be useful in constructing a more comprehensive view of NSC quiescence. NSC autofluorescence also provides a platform to identify nuanced substates. For example, in our *in vitro* NSC quiescence exit timelapse we were able to identify NSCs that were transitioning between quiescence and activation, as opposed to still quiescent or already fully activated. In our *in vivo* dataset, we find that PAF correlates with previously published substates of quiescence and activation. As this technology continues to develop, NSC autofluorescence could be used to further identify NSC substates both *in vitro* and *in vivo* as NSCs exit quiescence and differentiate. Importantly, we provide evidence that PAF intensity levels can be used to predict underlying gene expression and cell behavior. With this knowledge in hand, future studies will not require destroying the cell as with single cell sequencing, but can use this associated label-free signal as a readout of cell state identity and changes longitudinally to precisely map cell state and behavior throughout NSC quiescence exit and the entirety of adult neurogenesis more broadly.

Past using NSC autofluorescence to identify cell states, it could also be useful to understand what specifically is driving the changes in autofluorescence. We found that one of the most robust autofluorescent markers of NSC quiescence was high levels of PAF localizing to a subset of lysosomes. However, it remains unclear what specific autofluorescent molecule(s) is localized to lysosomes in qNSCs that causes them to be autofluorescent. The fact that there are dozens of well-known, and likely many unknown, highly autofluorescent molecules in the cell, combined with the broad spectral absorption and emission of qNSC lysosomes makes it challenging to identify what molecule is driving the autofluorescence [26]. Identifying the source of this signal could provide new insight into mechanisms underlying NSC quiescence. Importantly, many studies have found an increase in lysosomes in qNSCs [6, 13, 34]. Unique enzymes in lysosomes allow them to not only degrade proteins, but also lipids through endocytosis or lipophagy [46]. As qNSCs are performing fatty acid oxidation to provide energy for the cell [7] the lysosome’s ability to break down lipids and provide fatty acids for the cell, render them an important part of lipid metabolism in qNSCs. OCSI measures changes in lysosomal and mitochondrial metabolism, which act as a live and dynamic reporter of the underlying metabolic changes driving shifts in cell state.

Taken together, here we describe a single-cell resolution, non-destructive, graded, live-cell, label-free tool to study NSC quiescence with promise for the study of other types of stem cells and shifts in other types of NSC cell behavior, such as differentiation. We found that NSC autofluorescence, and specifically PAF intensity, was sufficient to predict NSC activation state in the absence of any exogenous labels. Finally, using this marker paired with single-cell transcriptomics, we define molecular mechanisms linked to rapid NSC activation and deep quiescence.

## Materials and Methods

### Mice

C57BL/6J mice were purchased from the Jackson Laboratory and used to establish a colony which was then used for experiments to isolate hippocampal NSCs for *in vitro* experiments. C57BL/6J mice from Envigo were used for the acute preparation from the SVZ, cell sorting, and sequencing experiments. Only males between 6 and 12 weeks of age were used for this study. All facilities used for maintaining the colony of mice used in this study have been approved by the Research Animal Resources and Compliance (RARC) at UW-Madison.

### Imaging Paradigms/Microscopes

#### 2-photon Imaging

Fluorescence lifetime images were captured on an Ultima (Bruker) two-photon microscope paired with a Nikon TiE body with a Chameleon Ultra II femtosecond-pulsed tunable Ti:Sapphire laser source (Coherent Inc., Santa Clara, CA, USA), a H7422PA-40 GaAsP photomultiplier tube (Hamamatsu Corporation, Bridgewater, NJ, USA) and a SPC-150 (Becker & Hickl) TCSPC card, using a Nikon CFI Apo LWD Lambda S 40XC water immersion objective. NAD(P)H was imaged by tuning the laser to 750 nm for two-photon excitation and collecting light using a 440/80 nm bandpass filter (Chroma). APC was imaged by tuning the laser to 890 nm for two-photon excitation and collecting light using a 550/100 nm bandpass filter (Chroma). mCherry was imaged by tuning the laser to 740 nm for two-photon excitation and collecting light using a 550/100 nm bandpass filter (Chroma). The field of view (256×256pixels) was scanned for 60 seconds with a 4.7 microsecond pixel dwell time.

#### Confocal Imaging

Confocal images were acquired using a Nikon C2 confocal microscope equipped with one fixed and one tunable GaAsP detector using excitation and emission described in the text. In general, autofluorescence images represent a single optical plane in the cell taken with 2-4x averaging. AP is ∼50-100x dimmer than conventional fluorophores, and thus, significantly higher powers are needed to visualize the signals described in this study. Specific excitation and emission parameters vary by experiment as listed in either Table 2 or each figure legend.

#### Fluorescent Activated Cell Sorting/Flow Cytometry

A BD FACSAria high speed cell sorter was used for all cell sorting experiments. A BD LSR II flow cytometer was used for analysis experiments. Specific excitation and emission parameters vary by experiment as listed in either Table 2 or each figure legend.

### NSC Dissection and Culturing

NSCs were isolated from the hippocampus by dissecting hippocampi from 3-5 male mice into cold HBSS, pooling samples, and then dissociating the tissue using GentleMACS Dissociator (Miltenyi Biotec) and MACS Neural Tissue Papain Dissociation Kit (Miltenyi Biotec 130-092-628) using the manufacturer’s protocol with added myelin removal, similar to previously described protocols [13, 47]. aNSCs were cultured as previously described at 37°C/5% CO2 in serum-free media (aNSC media): DMEM/F12 GlutaMax (Invitrogen 10565018) with B27 (1:50, Invitrogen 17504044), penicillin-streptomycin-fungizone (1:100, Invitrogen 15140122), and 20 ng/mL FGF-2 and EGF (PeproTech 100-18B and AF-100-15) [13, 47]. When culturing aNSCs as monolayers, cells were additionally treated with 5 μg/mL Heparin (Sigma H3149).

To image aNSCs as monolayers, aNSCs were plated onto glassware suitable for imaging (such as Fisher Scientific 12-565-337 or Ibidi 80826-G500) that was precoated with poly-L-ornithine for 1 hour at 37°C (PLO; 10 μg/mL plastic, 50 μg/mL glass, Sigma P3655) and laminin for 3 hours at 37°C (5 μg/mL, Sigma L2020). To create single cell suspensions for plating aNSCs, aNSCs were trypsinized using the following protocol: Cells were pelleted in a centrifuge at 120xg for 4 minutes and then treated with 0.05% trypsin (Invitrogen 25300-054) made in Versene (Thermo Fisher 15040066) at 37°C for 5 minutes [13]. Cells were then treated with twice the volume of trypsin inhibitor (Sigma T6522) for 2 minutes at room temperature, mechanically triturated and then pelleted again in a centrifuge by spinning at 120xg for 4 minutes. Single cells were then suspended and plated in aNSC media.

To generate qNSCs, aNSCs were plated onto PLO- and laminin-coated plates and treated with BMP-4, with FGF-2 and without EGF (qNSC media), using previously described protocols [6, 7, 13, 23, 24]. qNSC media is similar to aNSC media with the exception of the removal of EGF and the addition of 50 ng/mL BMP-4 (Fisher Scientific 5020BP010). After initial induction of quiescence, qNSCs were fed at least once every two days and were considered quiescent after 3 days of qNSC media treatment. For inducing quiescence exit, qNSCs were treated with aNSC media with the addition of 0.5 μg/mL of the BMP-4 antagonist noggin (PeproTech 120-10C) to more reproducibly induce quiescence exit.

### EdU Pulse

To measure the proliferation rate of NSCs, cells were pulsed with 10 μM EdU (Invitrogen C10337) to label cells progressing through S-phase of the cell cycle for 1 hour at 37°C. Following the pulse with EdU, cells were fixed in 4% paraformaldehyde (PFA) at room temperature (RT) for 15 minutes, and then treated to visualize EdU using a click-chemistry kit (Invitrogen C10337). To quantify EdU pulses, cells were costained with 20 μM Hoechst (Thermo Fisher Scientific 62249) to label nuclei of all cells. The total number of EdU+ and Hoechst+ cells were then counted across at least three experiments performed on different days and analyzed per image.

### Analysis of Fluorescence Lifetime Imaging Data

Fluorescent lifetime images were analyzed using SPCImage version 8.3. For each fluorescent lifetime image, background was removed through thresholding (value of 50 for NAD(P)H, value of 5 for APC). Fluorescence lifetime decays were deconvolved from the instrument response function and modeled by fitting to a biexponential decay equation: I(t) = α_1_*e^-t/τ1^ + α_2_*e^-t/τ2^ + C, when I(t) is the autofluorescent intensity as a function of time (t) after the laser pulse, α_1_ and α_2_ are fractional contributions (α_1_ + α_2_ = 1) of long and short lifetime components, τ_1_ and τ_2_ reflect the short and long lifetime components and C represents background photons. The fluorescence lifetime (T_m_) was defined as: T_m_ = α_1_*τ_1_ + α_2_*τ_2_. After fitting the data, average fluorescence lifetime and intensities were calculated per cell examining only the cytoplasm. To identify cytoplasm, CellProfiler was used to generate masks around the entire cell and the cell’s nucleus and then nuclear space was subtracted from the whole cell to identify only cytoplasmic signal. Average endpoint values for each cell, were obtained by using R to calculate average values using the CellProfiler generated cytoplasm masks.

### Dimension Reduction Analyses

PCA was used to reduce the dimensions of NSC autofluorescence imaging data using the PCA() function in R. To perform the PCA, non-redundant NSC autofluorescence endpoints (intensity, α_1_, τ_1_, and τ_2_) were normalized to values within 0 and 1 and used for the analysis.

### Machine Learning Analyses

Random Forest models were developed to assess the capacity of NSC autofluorescence to classify NSC activation state using the randomForest() function in R and used to generate the representative ROC plots shown in this manuscript [48]. Non-redundant NSC autofluorescence endpoints (intensity, α_1_, τ_1_, and τ_2_) were normalized to values within 0 and 1 and then varying subsets of these endpoints, as described in the text, were used to generate each Random Forest model.

### Autofluorescence Imaging with Organelle Dyes

To image NSC autofluorescence with various markers of other organelles in the cell, qNSCs and aNSCs were generated as described above and then treated with the following dyes and specifications. Lysotracker Deep Red (Thermo Fisher Scientific L12492) was added to NSCs at a dilution of 1:7000 and incubated on qNSCs for 10 minutes in the incubator prior to imaging. To avoid bleed-through from the dyes, autofluorescence was imaged by exciting with a 405 nm laser and collecting at 525-560 nm.

### Flow Cytometry NSC Cell State Enrichment Assays

A BD FACSAria high speed cell sorter was used to sort NSCs. qNSCs and aNSCs were pulsed with EdU for one hour as described above in the incubator and then put into single cell suspension – aNSCs by trypsinizing and qNSCs by just mechanically lifting from the dish. Cells were counted and then a 1:1 mix of aNSCs and qNSCs was made. All samples were put in phosphate buffered saline on ice until completion of the sort. On the cytometer, forward and side scatter was used to identify single cells and then NSCs were sorted by autofluorescence using a 405 nm laser to excite and collecting light between 565 and 645 nm. Gates were drawn to collect the brighter or dimmer NSCs and then samples were collected and plated onto glassware coated in PLO and laminin. 2-3 hours after plating following cell sorting (after cells adhered to the dish) samples were fixed in 4% PFA for 15 minutes at room temperature. Cells were then stained and analyzed for EdU as described above.

### Quiescence Exit Timelapse

qNSCs were generated and then sequentially treated with activation media in such a way that qNSCs that had been exiting quiescence for 0 hours, 24 hours, 48 hours, 72 hours and aNSCs could be imaged at the same time. 2-photon FLIM autofluorescence imaging was performed and analyzed as discussed above. To perform the quiescence exit EdU pulse, samples were prepared for each step in the time course and then treated, processed and analyzed as discussed in the “EdU Pulse” section of the methods.

To label aggregated proteins with Proteostat (Enzo ENZ-51035-0025), NSCs were prepared for each part of the time course and then fixed in 4% PFA for 15 minutes at room temperature. Cells were stained in accordance with the manufacturer’s protocol. In brief, cells were permeabilized with 0.5% triton in 0.5 mM ethylenediaminetetraacetic acid (EDTA) in PBS for 30 minutes at room temperature and stained with Hoechst (1:5,000) to visualize nuclei and Proteostat (1:1,000) for 30 minutes at room temperature. Following staining with the dyes, cells were washed 3 times in PBS for 10 minutes at room temperature. To analyze Proteostat signal, number of Proteostat puncta were counted per cell over at least 5 images per condition. The experiment was repeated for a total of 3 times.

### AAV4 Particle Production

AAV4 particles were produced by the Duke Viral Vector Core. The AAV4 vector was a generous gift from Dr. Juan Song.

### BrdU Labeling

Mice were injected with AAV4-CMV-mCherry particles and then allowed to express for 3 days, mimicking a previously described protocol [37]. On the third day following AAV4 injection, BrdU was dissolved 0.02 g/mL in PBS and then administered to mice by intraperitoneal injection at 200 mg/kg, a total of 3 times over the course of 4 hours (each injection 2 hours apart). 2 hours following the final BrdU injection, mice were transcardially perfused with 0.9% saline followed by 4% PFA in a 0.2M phosphate buffer. Brains were then extracted and postfixed in 4% PFA overnight at 4°C and then stored in 30% sucrose in phosphate buffered saline until the brains sank (∼1-3 days). Brains were then sectioned on a sliding microtome to generate 40 μm thick slices and then immunostained for the following markers: BrdU (Abcam ab6326), mCherry (Clontech 632496; to amplify mCherry signal) and 4′,6-diamidino-2-phenylindole (DAPI; Invitrogen D1306; to label nuclei) using the following protocol. Sections were rinsed 3 times for 10 minutes in tris buffered saline (TBS) at room temperature and then incubated with 1 M HCl for 20 minutes at 37°C, incubated in 0.1 M Borate Buffer 2 times for 15 minutes at room temperature, rinsed 3 times with TBS for 10 minutes each at room temperature, blocked in 0.25% TritonX-100, 3% Donkey Serum (Millipore Sigma S30-100ML) in TBS (TBS++ buffer) for 1 hour at room temperature, incubated with primary antibodies overnight at a dilution of 1:500 in TBS++ at 4°C, rinsed 3 times in TBS at room temperature for 10 minutes, incubated with secondary antibodies 1:500 for 1.5 hours at room temperature in TBS++, incubated with DAPI (1:1,000 in TBS) for 5 minutes at room temperature, and then rinsed 2 times in TBS at room temperature prior to mounting slides.

Samples were then imaged and analyzed to determine whether mCherry+ cells were BrdU+. Representative series of brain sections (∼8 slices per series) were analyzed over a total of 3 animals. All mCherry+ cells in the dentate gyrus were analyzed.

### Stereotaxic Injections and Organotypic Slice Culture Experiment

To inject AAV4 particles into the dentate gyrus of the mouse brain, we adapted previously reported protocols [37, 44]. Mice were anesthetized with isoflurane, and then mounted in a stereotaxic frame (Stoelting). Optixcare Pet Eye Lubricant (Amazon B0799LS6ZY) was applied periodically to maintain eye lubrication. The incision site was sterilized with Betadine and then cut to access the skull. The incision site was then anesthetized with a splash block of 50:50 lidocaine/bupivacaine. Following local anesthetic, two holes were drilled manually with a needle over the dentate gyrus at coordinates: AP −2.0 mm, ML +/-1.4 mm. A Hamilton syringe (Sigma-Aldrich 20734) loaded with AAV4 particles was then inserted into each hole (one at a time) and inserted down 2.8 mm ventral to the cortical surface. After inserting the needle, the needle was left for 2 minutes, and then 1 uL of AAV4 (∼1*10^8 particles/uL) was injected over the course of 7 minutes. Following injection of the virus, the needle was left for an additional two minutes prior to removing the needle and proceeding. Following removal of the needle, the skin was moistened with 0.9% saline, and then resealed with a chemical suture (3M Vetbond Tissue Adhesive). Animals were administered 5mg/kg Meloxicam through IP injection as a preemptive anesthetic following surgery. Following surgeries, animals were monitored until they woke up and then administered additional meloxicam in the days following surgery as necessary if any signs of discomfort were observed.

To generate slice cultures, we adapted a previously published protocol [38]. 7 days following injection (viral expression was not detectable in live slices until ∼7 days after injection) mice were euthanized and brains were extracted and immersed in ice-cold 70% ethanol for 2 minutes. Following ethanol treatment, brains were mounted onto the stage with superglue and then sectioned at 400 μm thick on a Leica VT1000S vibratome in ice-cold dissection media (95.2 g/L Hank’s Balanced salt (Sigma H2387), 4.2 mM NaHCO_3_, 10 mM HEPES (Thermo Scientific 15630106), 33.3 mM Glucose, 1X Penicillin/Streptomycin (Invitrogen 15140122), 0.3% BSA (Sigma-Aldrich A2153-50G), 12 mM MgSO_4_-7H_2_O). Following slice generation, slices were placed in culturing medium (50% MEM (Invitrogen 11575032, 25% Hank’s Balanced Salt (Invitrogen 24020117), 25% Horse Serum (Invitrogen 26050070, 12.5 mM HEPES, 1X Penicillin/Streptomycin (Thermo Scientific 15630106), 35 mM Glucose), placed in a 37°C/5%CO_2_ incubator, and imaged within 3 hours.

### Analysis of autofluorescence intensity/area, particle number and area covered, and colocalization

To obtain the autofluorescence intensity/area, particle number and area, as well as colocalization of signals, at least three images were taken per condition in one plane using a confocal microscope, and subsequently analyzed using ImageJ. In short, brightness and thresholds were established for each channel, followed by “Despeckle” to reduce background noise. Each image was separately quantified for particle number and area covered using “Analyze Particles” (Size (micron^2)=0.08-40; Circularity=0-1). To detect colocalization of particles, ROI overlays for detected particles of Lysotracker were placed on the autofluorescence (AF) image. AF within ROIs was considered colocalized. AF particles outside of these ROIs were further studied to determine if they 1) were not colocalized, or 2) were within 2microns of a ROI with no particle present, suggesting the particle moved between images, thus still considered colocalized.

*(AF total particle number – Particles outside of Lyso ROIs) / Total AF particle number = Percent of AF colocalized with signal Number of colocalized AF particles / Total Lyso particles = Percent of Lysotracker or Lipidspot signal colocalized with AF*

### *In vitro* SORT-seq

Primary mouse hippocampal NSCs were cultured in full media and plated into 4 wells of a PLO-, laminin-coated 6-well plate. At separate time points, cells were induced into quiescence. For quiescence exit conditions, following 3 days of quiescence media, the media was changed to activation media. All conditions were prepared so that they would be ready for sorting at the same time. Activated NSCs were plated separately 2 days before collection. The day of the sort, NSCs were trypsinized from the dish (0.25% Trypsin-EDTA) for 3 minutes, followed by quenching with trypsin inhibitor. Following a spin of 120xg for 4 minutes, cells were resuspended in ice-cold FACS buffer (sterile-filtered, ice cold DPBS, 1mM EDTA, 4ug/ml DNAse), and taken immediately to the cell sorter. Sort-seq plates were thawed on ice at least 15 minutes prior to collection, and at the sorter the collection device was kept at 4 degrees C. 47 cells/gate were collected for each of 4 gates (188 cells total/cell state equally distributed) for each condition. Plates were spun down, snap frozen, and sent for library preparation and sequencing to Single Cell Discoveries.

### *In vivo* SORT-seq

This experiment was performed 3 separate times, collecting cells from each experiment to be analyzed together. 8 male Envigo C57BL/6J mice per experiment were used to sort cells from the SVZ. Using a modified previously published protocol for dissection [39], SVZ tissue was dissected into cold HBSS without Ca2+ and Mg2+, followed by mincing of the tissue with a razor blade. Tissue was resuspended in 0.05% Trypsin-EDTA, and put in a 37 degrees C water bath for 6 minutes. Trypsin was quenched using a double volume of pre-warmed trypsin inhibitor including DNAse, and centrifuged for 300xg for 5 minutes. The pellet was resuspended in a cold, sterile-filtered, 0.5% BSA/PBS solution (sort buffer), and dissociated by pipetting 20 times up/down. A 45μM mesh filter was pre-wet with the same solution, and the run-through discarded. The brain sample was run through the filter, followed by a 1ml rinse. The cell solution was spun for 300xg for 5 minutes for 4 degrees C. All spins and treatments from this point forward were performed at 4 degrees C. For 2 of the 3 experiments, myelin was removed using a Miltenyi Magnetic sorting protocol. In short, the appropriate volume of cold sort buffer was added based on weight, as well as myelin removal beads, followed by incubation for 15 minutes in the refrigerator. After washing, lysate was centrifuged at 300xg for 10 minutes. The pellet was resuspended in sort buffer, and then was filtered through the magnetic column. The effluent was spun down, and then resuspended in sort buffer and an anti-GLAST-Biotin antibody (Miltenyi), incubating for 10 minutes in the refrigerator. Following wash and centrifugation, cells were further labeled with anti-biotin microbeads (Miltenyi) for 15 minutes in the refrigerator. To prepare cells for loading onto the magnetic column, they were washed, centrifuged, and resuspended in sort buffer. Cells that were GLAST+ were on beads that remained in the column on the magnetic plate. The column was washed, and beads and cells eluted into a collection tube using a plunger once removed from the magnetic plate, centrifuged, and resuspended in an ice-cold staining buffer [40] containing L-15, 1M HEPES 7.4 pH, 100x Pen-strep, and 100mM EDTA. CD24-APC-eFluor780 (M16947-0242-82; 1:200) and O4-Cy7 (Miltenyi – 130-115-810; 1:200) were added for a total of 45 minutes, with CD133-APC (Clone 13A6; 1:50) added in the final 25 minutes of incubation on ice. Cells were then washed and centrifuged, and resuspended in FACS buffer with 4μg/ml DNAse, and transported to the FACS machine on ice. There, using previously established compensated settings, GLAST+ (per bead sorting) CD133+ (low to middle intensity) CD24-, O4-(together labeled in the same “dump” channel) were sorted into a 384 well plate, with autofluorescence intensity measured for each cell. To establish the compensated settings, cells collected from the brain were singly labeled for each antibody, and then as a group.

Viable single cells were FACS sorted into 384-well plates, called cell capture plates, that were ordered from Single Cell Discoveries, a single-cell sequencing service provider based in the Netherlands. Each well of a cell capture plate contains a small 50 nl droplet of barcoded primers and 10 μl of mineral oil (Sigma M8410). After sorting, plates were immediately spun and placed on dry ice. Plates were stored at −80°C.

Plates were shipped on dry ice to Single Cell Discoveries, where single-cell RNA sequencing was performed according to an adapted version of the SORT-seq protocol [35] with primers from [49]. Cells were heat-lysed at 65°C followed by cDNA synthesis. After second-strand cDNA synthesis, all the barcoded material from one plate was pooled into one library and amplified using in vitro transcription (IVT). Following amplification, library preparation was done following the CEL-Seq2 protocol [50] to prepare a cDNA library for sequencing using TruSeq small RNA primers (Illumina). The DNA library was paired-end sequenced on an Illumina Nextseq™ 500, high output, with a 1×75 bp Illumina kit (read 1: 26 cycles, index read: 6 cycles, read 2: 60 cycles).

During sequencing, Read 1 was assigned 26 base pairs and was used for identification of the Illumina library barcode, cell barcode and UMI. R2 was assigned 60 base pairs and used to map to the reference transcriptome of Hg19 with BWA [51]. Data was demultiplexed as described in Grün et al. 2014. Mapping and generation of count tables was automated using the MapAndGo script.

### SORT-seq Analysis

The data from all samples were loaded in R (version 4.2.0) and processed using the Seurat package (version 4.1.1 [52]). More specifically, cells with at least 5000 UMI’s per cell and less than 15% mitochondrial gene content were retained for analysis of the *in vitro* dataset. Cells with at least 1000 UMI’s per cell and less than 10% mitochondrial gene content were retained for analysis of the *in vivo* dataset. The datasets were normalized for sequencing depth per cell and log-transformed using a scaling factor of 10,000 and used for dimensionality reduction and clustering. Cells were clustered using graph-based clustering and the original Louvain algorithm was utilized for modularity optimization. The differentially expressed genes per cluster were calculated using the Wilcoxon rank sum test and used to identify cell types.

Cell cycle was scored using the CellCycleScoring function of Seurat, adjusted to include G0 genes for the scoring of a cell cycle. A list of ten G0-specific genes (Scrg1, Plp1, S100b, Gpm6b, Bex1, Ptprz1, Prcp, Ptn, Sox4, Sat1) were retrieved from [36].

Fluorescence levels were recorded during index sorting of the single cells in cell capture plates. Autofluorescence levels were added as metadata for each cell. To infer the difference between autofluorescence levels, differential expression analysis was performed using the Wilcoxon test implemented in Seurat’s FindMarkers function. Cells from each condition were ranked and binned based on their autofluorescence levels. The cells with autofluorescence levels falling in the top 15% (*in vitro* dataset) or 30% (*in vivo* dataset) were compared to the bottom 15% or 30% cells. Regardless of the statistical significance, the top genes are represented in a heatmap showing the expression variation across cells with different autofluorescence levels.

The gene list resulting from the differential expression test were subjected to gene set enrichment analysis. Unfiltered gene lists were ranked based on fold change and used as input for the enrichment test using fgseaSimple (minSize=15, maxSize=600, nperm=10000) implement by the fgsea package (version 1.22.0). The gene lists were tested for enrichment against the mouse hallmark gene sets/curated gene sets/ontology gen sets collection of molecular Signatures Database [53, 54]. GO term analyses were performed using GOrilla (http://cbl-gorilla.cs.technion.ac.il/), reporting “All” ontology terms using gene lists that were differentially expressed at a log2(fold change) of 0.75 or greater in each analysis respectively. PAF Intensity compared against expression of genes are shown with cells with a value of “0” for expression removed.

To compare our analysis with Llorens-Bobadilla et al 2015, a counts file was generated from combining cells labelled as aNSC1, qNSC1, aNSC2 or qNSC2 in GSE67833 [10] and cells from clusters 3 and 4 in our *in vivo* data set. Using custom Python scripts, the counts file was filtered to exclude cells with less than 500, or more than 9000 genes. Any gene detected in 1 cell or more was kept. UMAP embedding was performed using the Python Scanpy package (PMID: 29409532) with min_dist=0.25. Autofluorescence was presented as a heat map based using the default Plotly continuous color palette.

### Statistics and replicates

Experiments, unless otherwise noted, were replicated on at least three separate days with at least three technical replicates each day as applicable. Significance and analyses were performed in either R Studio, Microsoft Excel, or GraphPad Prism using tests indicated in the figure legends. To analyze 2-photon imaging data (FLIM data), a Generalized Linear Model was formed to assess significance in R Studio. All other statistical tests were performed in GraphPad Prism.

### Data Accessibility/Code

In addition to supplementary data files attached to this manuscript, all code and data will be available upon reasonable request to the corresponding author: Dr. Darcie L. Moore. All SORT-seq data sets are available at the gene expression omnibus (GEO): X.

## Acknowledgments

We thank the UW-Madison flow cytometry core (P30 CA014520 and 1S10RR025483-01); J. Song for the AAV4 construct; and members of the Moore lab and UW-Madison community for their input. We thank BioRender for supplying graphics used to make schematics (BioRender.com). We thank Single Cell Discoveries for their help with project design, single-cell sequencing services, and data analysis. We thank our funding sources: NIH T32 T32GM008688 (to C.S.M.), Diana Jacobs Kalman Fellowship from AFAR (to C.S.M.), Wisconsin Graduate Fellowship (to C.S.M.), DP2 NIH New Innovator Award (to D.L.M.), and Vallee Scholar Award (to D.L.M.).

## Author Contributions

C.S.M., D.L.M., and M.C.S. conceived the idea and planned the study. C.S.M. performed most of the experiments. K.T. and A.J.W. assisted with the *in vitro* FLIM experiments. Z.P.A. assisted with validation of AAV labeling and quantified colocalization of puncta with other markers. B.P. performed the Lysotracker colocalization experiment and Proteostat time course. R.D.R. performed the vimentin expression timelapse. M.M.C. performed Proteostat analyses. E.P.W. and M.V.J. assisted in establishing a slice culture protocol. P.C.K. assisted with several *in vitro* experiments and the *in vivo* single cell experiment. D.L.M. generated the *in vitro* and *in vivo* samples for single-cell RNA sequencing. A.R. performed the comparative transcriptomics analysis. C.S.M. and D.L.M. wrote the manuscript with input from all authors.

## Supplementary Table Legends

Table S1 – Transcript counts from *in vitro* SORT-Seq

Table S2 – Differential expression analyses from *in vitro* SORT-Seq comparing the top 15% PAF intensity cells to the bottom 15% PAF intensity cells in each condition separately

Table S3 – Enriched gene ontology terms from the differential expression analysis of the top 15% PAF intensity cells to the bottom 15% PAF intensity qEx 48 hr transcriptomes from *in vitro* SORT-Seq

Table S4 – Transcript counts from *in vivo* SORT-Seq

Table S5 – Differential expression analyses of the top third compared to the bottom third PAF intensity cluster 3 qNSCs from *in vivo* SORT-Seq

Table S6 – Enriched gene ontology terms from the differential expression analysis of the top third compared to the bottom third PAF intensity cluster 3 qNSCs from *in vivo* SORT-Seq

**Figure S1.**
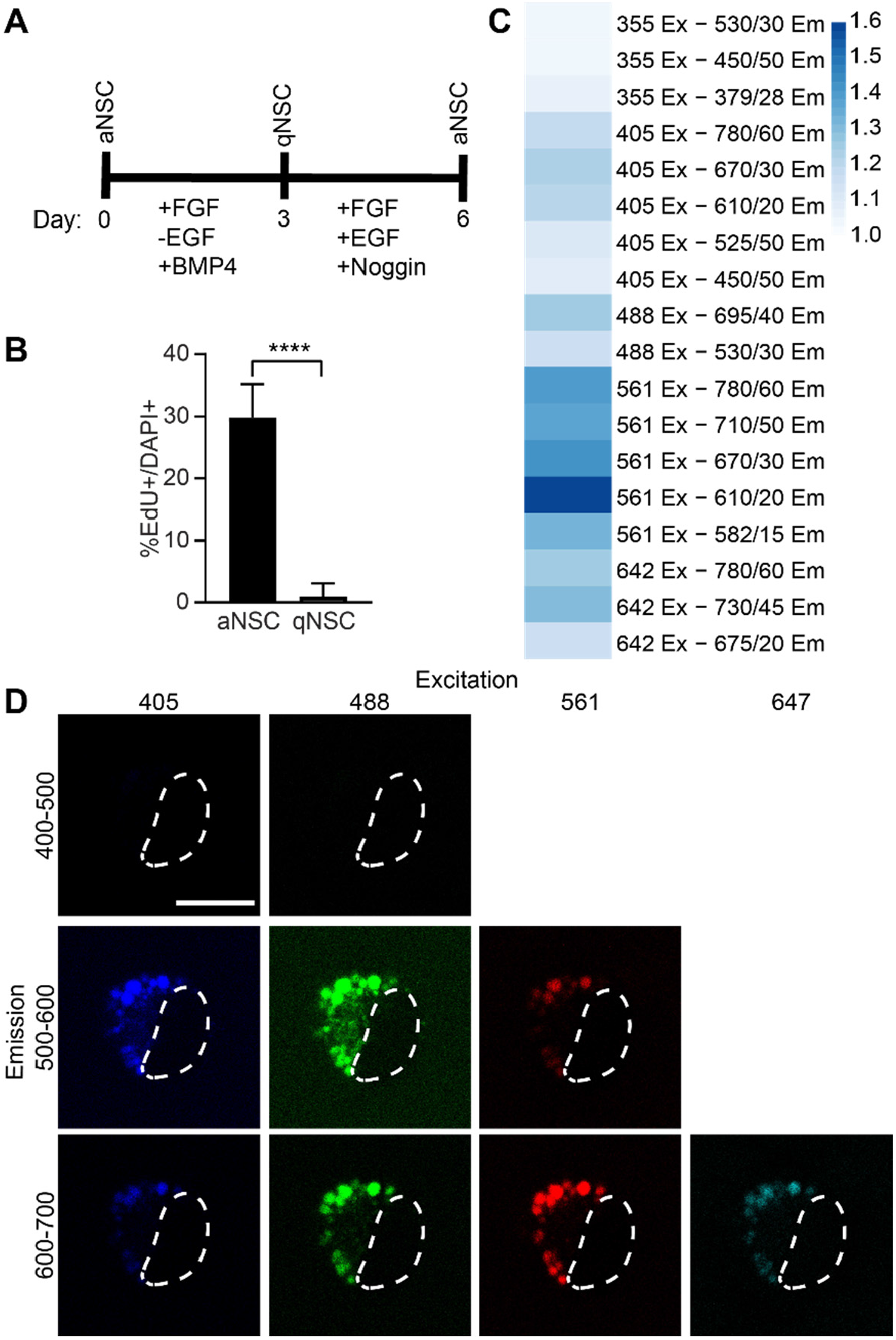
A) Schematic of the BMP4 system for inducing reversible NSC quiescence. B) aNSCs and qNSCs were pulsed with EdU for 1 hour and then fixed, stained with DAPI, and analyzed for %EdU+ cells over total DAPI+ nuclei (N=3, Student’s t test, mean ± SD). C) Heat map depicting differences in qNSC autofluorescence relative to aNSC autofluorescence measured on a flow cytometer. D) qNSCs were imaged using a confocal microscope with excitation and emission as specified in the figure. Scale bar, 10 μm. ****p < 0.001.

**Figure S2.**
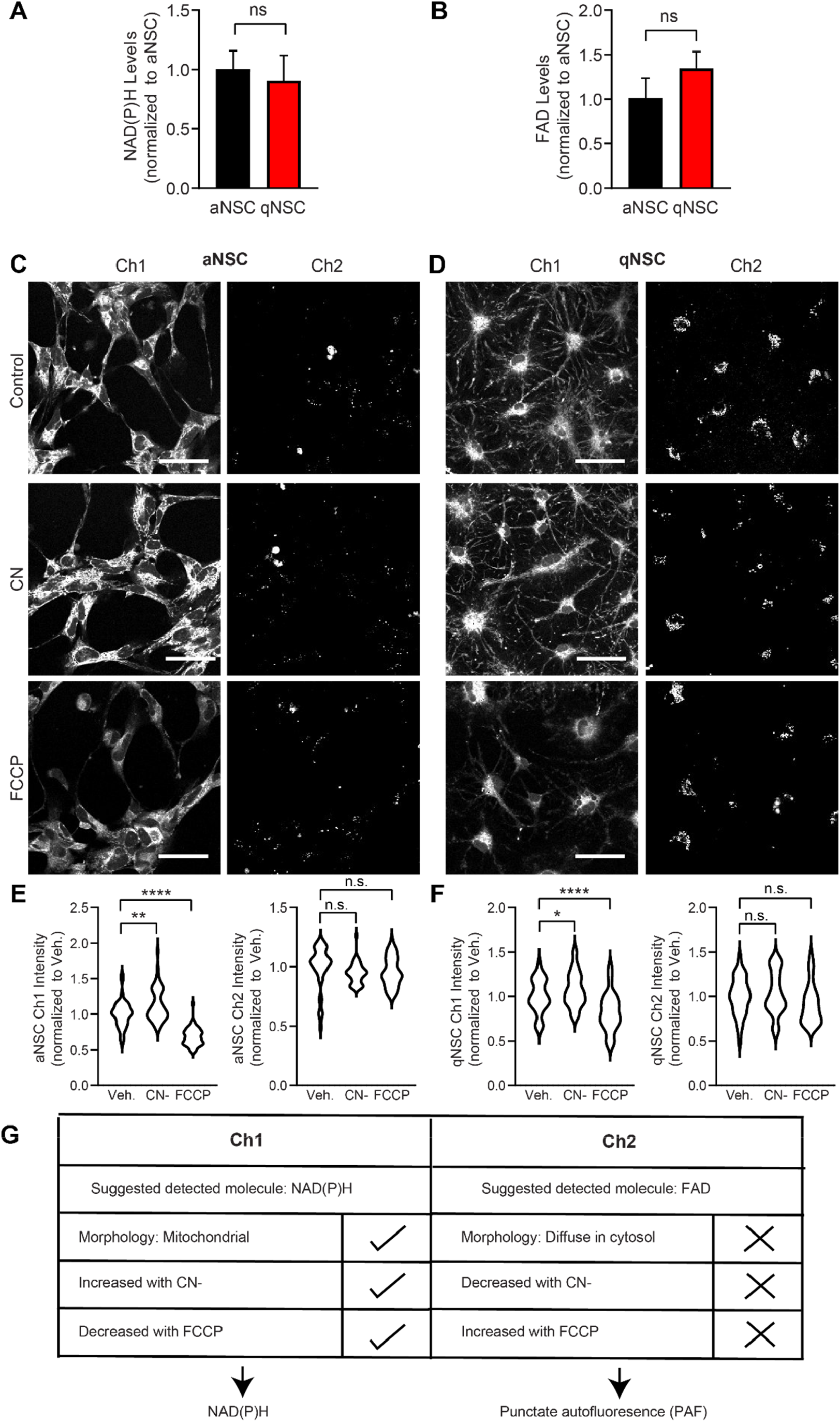
A-B) aNSCs (black bars) and qNSCs (red bars) were lysed and analyzed for relative levels of NAD(P)H and FAD (N=3, Student’s t test, mean ± SD). C-F) qNSCs were either untreated (control), treated with 2 mM CN^-^ (red bar), 1 μM FCCP (blue bar), or 1% DMSO vehicle control for 20 minutes and then analyzed for Ch1 and Ch2 intensity (N=3, Student’s t test, mean ± SD). G) A summary of our fundings from Fig S2 results in further discussion of Ch1 as NAD(P)H, and for Ch2, PAF. Scale bars, 50 μm. *p < 0.05, **p < 0.01, ****p < 0.0001.

**Figure S3.**
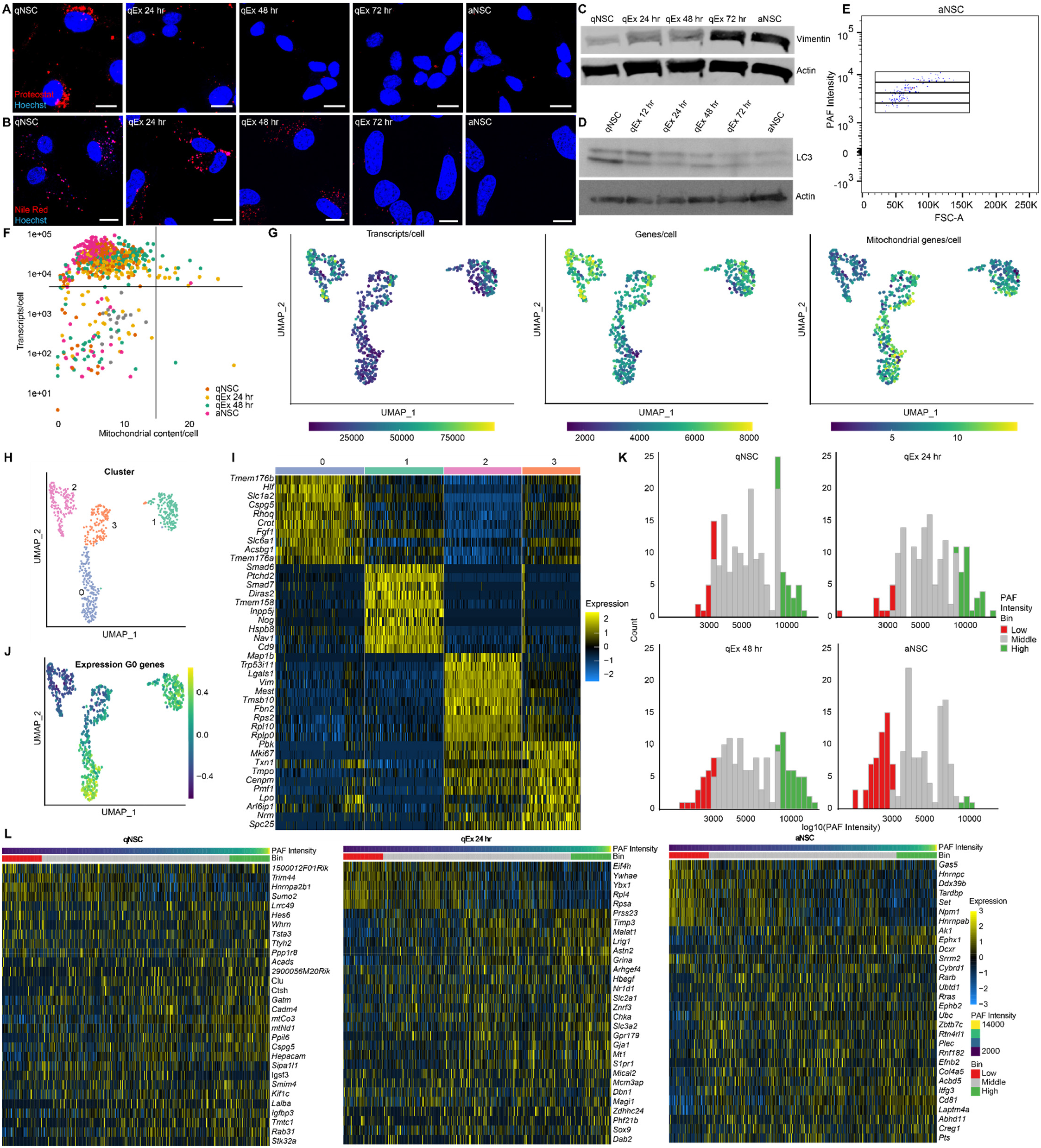
A) Representative Proteostat images of the data shown in Fig. 3I. NSCs during the fixed quiescence exit time course were fixed and stained to visualize aggregated proteins (Proteostat; red) and nuclei (Hoechst; blue). B) Representative Nile red images of the data shown in Fig. 3I. NSCs during the fixed quiescence exit time course were labeled with Nile red (lipid droplets; red) and Hoechst (nuclei; blue). C) Western blot showing changes in vimentin expression during quiescence exit corresponding to the data shown in Fig. 3I. NSCs during the fixed quiescence exit time course were processed to extract total protein and vimentin and actin expression were probed by western blot. D) Western blot showing changes in LC3I and LC3II expression during quiescence exit corresponding to the data shown in Fig. 3I. NSCs during the fixed quiescence exit time course were processed to extract total protein and vimentin and actin expression were probed by western blot. E) FACS plot showing aNSCs analyzed by FACS and example bins used to isolate a broad range of PAF intensity for single cell sequencing. F) A plot of *in vitro* SORT-seq mitochondrial content per cell vs transcripts per cell. G) UMAP analysis of *in vitro* SORT-seq transcriptomes labeled by transcripts per cell, genes per cell and mitochondrial genes per cell shown. H) UMAP analysis of *in vitro* SORT-seq transcriptomes labeled by clusters identified using the Louvain method. I) Heat map showing normalized expression of the top 10 most differentially expressed genes marking each cluster identified through *in vitro* SORT-seq. J) UMAP analysis of *in vitro* SORT-seq cell transcriptomes labeled by expression of G_0_ genes [36]. K) Histograms illustrating bins used across conditions for differential expression analysis of high and low PAF intensity cells from *in vitro* SORT-seq. L) Heat maps showing normalized expression of the top 30 most differentially expressed genes distinguishing high from low PAF intensity qNSCs, qNSCs exiting quiescence for 24 hours and aNSCs from *in vitro* SORT-seq. Scale bars, 10 μm.

**Figure S4.**
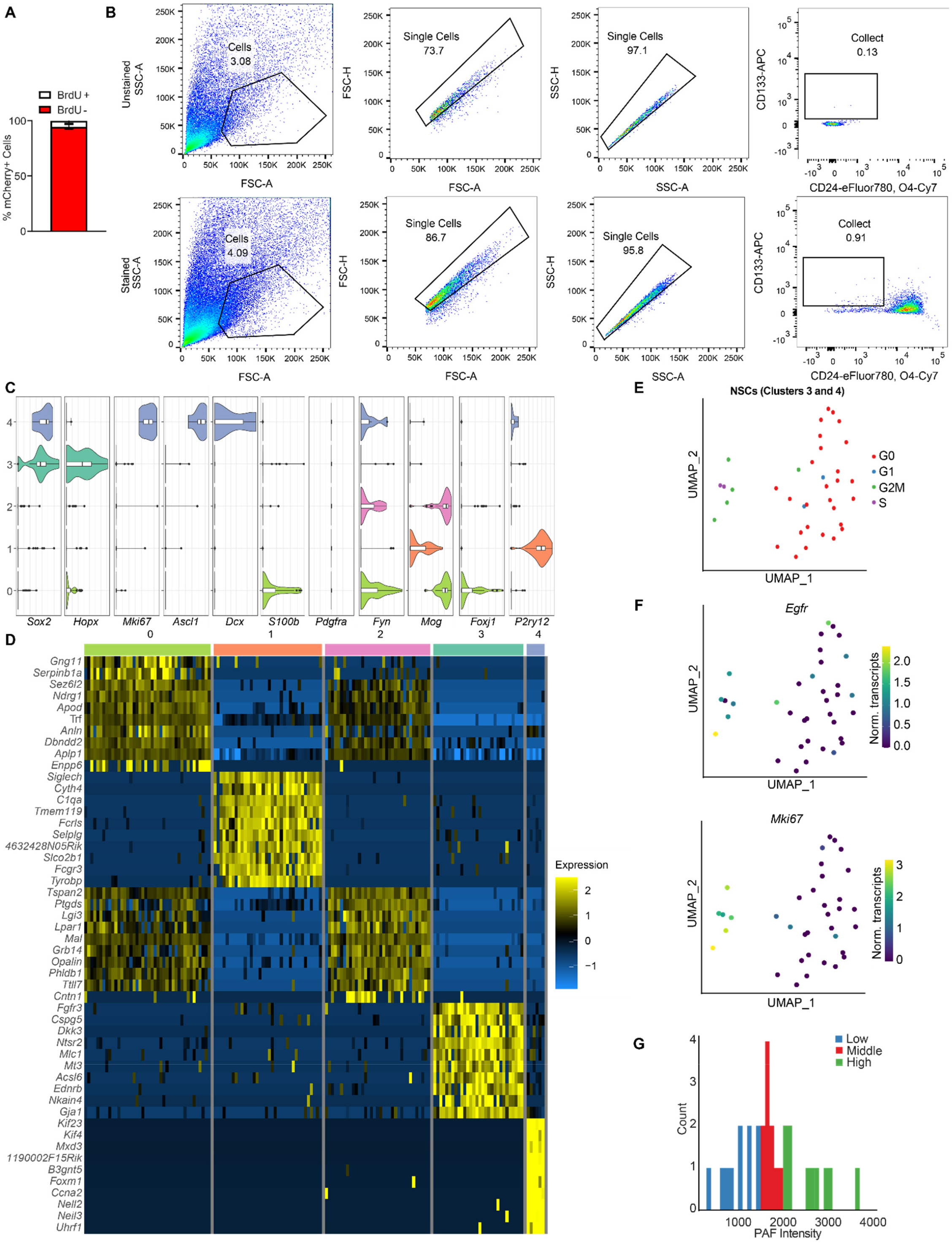
A) Mice were stereotactically injected with AAV4-CMV-mCherry viral particles into the dentate gyrus of the hippocampus, and allowed to recover for 3 days before receiving three BrdU injections (200 mg/kg) two hours apart prior to perfusion. Brains were extracted, sectioned, and immunostained to visualize BrdU and mCherry. AAV4-CMV-mCherry+ cells were counted as BrdU+ (white bar) or BrdU-(red bar) (n=3, mean ± SD). B) FACS strategy for enriching for NSCs from the SVZ. C) Violin plots depicting expression of marker genes across clusters identified from SORT-seq of NSC-enriched SVZ samples. D) Heat map showing normalized expression of the top 10 most differentially expressed genes marking each cluster identified through *in vivo* SORT-seq. E) UMAP analysis of cluster 3 and 4 *in vivo* SORT-seq transcriptomes labeled by predicted cell cycle stage. F) UMAP analysis of cluster 3 and 4 *in vivo* SORT-seq transcriptomes labeled by expression of *Egfr* and *Mki67*. G) Histogram depicting bins of PAF intensity across qNSCs (cluster 3) used for the differential gene expression analysis shown in Fig. 4K-L.

